# Transcriptome dynamics predict thermotolerance in *Caenorhabditis elegans*

**DOI:** 10.1101/661652

**Authors:** Katharina Jovic, Jacopo Grilli, Mark G. Sterken, Basten L. Snoek, Joost A. G. Riksen, Stefano Allesina, Jan E. Kammenga

## Abstract

**Background:** The detrimental effects of a short bout of stress can persist, and potentially turn lethal, long after the return to normal conditions. Thermotolerance, which is the capacity of an organism to withstand relatively extreme temperatures, is influenced by the response during stress exposure, as well as the recovery process afterwards. While heat-shock response mechanisms have been studied intensively, predicting thermal tolerance remains a challenge.

**Results:** Here, we use the nematode *Caenorhabditis elegans* to measure transcriptional resilience to heat stress and predict thermotolerance. Using high dimensionality reduction techniques in combination with genome-wide gene expression profiles collected in three high resolution time-series during control, heat stress and recovery conditions, we infer a quantitative scale capturing the extent of stress-induced transcriptome dynamics in a single value. This scale provides a basis for evaluating transcriptome resilience, defined here as the ability to depart from stress-expression dynamics during recovery. Independent replication across multiple highly divergent genotypes reveals that the transcriptional resilience parameter measured after a spike in temperature is quantitatively linked to long-term survival after heat stress.

**Conclusion:** Our findings imply that thermotolerance is an intrinsic property that pre-determines long term outcome of stress and can be predicted by the transcriptional resilience parameter. Inferring the transcriptional resilience parameters of higher organisms could aid in evaluating rehabilitation strategies after stresses such as disease and trauma.

## Background

Temperature is a key factor that directly affects physiological processes, life history and behaviour of many organisms. Ambient temperatures can rise suddenly, inflicting physiological consequences often lasting far beyond the initial exposure. For instance, it has repeatedly been shown that exposure to heat stress early in life can have an effect on traits later in life such as reproductive success and lifespan in the nematode *Caenorhabditis elegans* and fruit fly *Drosophila melanogaster* [1–5]. The ability to withstand the negative effects of heat stress is called thermotolerance and requires instant regulatory protective responses involving the well-studied heat-shock response [6]. Since tolerance is a trait that results in the absence of adverse effects, it is difficult to predict tolerance levels of an organism before the negative effects of stress have become apparent.

Next to the induction of genes within specific stress-response pathways, recent studies in *C. elegans* have shown that heat stress also induces a broad acclimation of transcriptional patterns involving differential expression of thousands of genes [7–9]. Furthermore, during prolonged stress exposure, expression changed continuously until lethal stress levels were reached [8]. Those findings illustrate that the state of the transcriptome directly reflects the stress levels the organism was exposed to. While the reactive processes occurring during the heat-shock response are well understood, much less is clear about how organisms recover from a heat shock and how the genome wide transcriptional state might be used to predict long-term outcome of a short bout of heat stress.

Here, we quantify gene expression resilience during and after heat stress in order to predict thermotolerance in *C. elegans.* First, by measuring genome-wide gene expression levels of the canonical laboratory strain Bristol N2 in three high-resolution time series (development, heat stress, and recovery from heat stress), and applying high dimensionality reduction techniques to the data, we show that the state of the transcriptome during and after the dynamic response to heat-stress perturbations can be captured by a single parameter. This finding provides the basis to evaluate and compare complex transcriptional patterns after stress in a straight forward and quantitative way.

Secondly, in order to generalize our findings beyond the individual genotype, we expanded our analyses across different genetic backgrounds. Previous research shows that different genotypes are differently affected by the heat stress [3, 9, 10], assumingly due to an intrinsic difference in thermal tolerance. Our results show that transcriptome resilience measured after a mild heat stress early in the development of *C. elegans* is predictive of its thermotolerance. Thermotolerance (based on long-term survival) and transcriptional resilience were measured in independent populations of the same genotype, emphasising the genetically intrinsic nature of thermal tolerance and the robustness of this approach to predict thermal tolerance. Our methods are straightforward to implement and allow to map gene-expression data during and after heat stress onto a few main quantitative scales that have a clear biological interpretation.

## Results and discussion

### Using dimensionality-reduction techniques to infer a developmental axis *D* and heat-stress axis *H*

*C. elegans* develops relatively fast – within ∼65 hours an individual develops from an egg into a reproducing adult [11]. The transition through the four larval stages is controlled by highly dynamic transcriptional processes [12, 13]. To characterize the temporal dynamics in genome-wide gene expression during heat stress and in recovering *C. elegans* populations, we have to remove stress-independent variation in gene-expression patterns caused by differences in development between samples collected in a time-series spanning several hours. For this purpose, we compiled a data set of 71 gene expression profiles measured in isogenic populations of the canonical strain Bristol (N2) sampled in an approximately hourly interval during exposure to three different treatments: i) during unperturbed development at 20°C [12], ii) during prolonged exposure to heat-stress conditions at 35°C [8], or iii) during a period of recovery at 20°C after a 2-hour heat stress at 35°C (Fig. 1A). First, the data was separated into training and testing sets. Second, through the application of dimensionality-reduction techniques (*i.e.* principal component analysis) on expression profiles from unperturbed populations (n = 9 samples out of 18) and from heat-stressed populations (n = 9 samples out of a total 39), we inferred the combination of gene-expression patterns that best characterized the overall expression dynamics of each treatment (Fig. 1B, Methods, and SI). From the unperturbed data, we obtained a *developmental axis D* (see SI section S2), describing the temporal expression dynamics during development. Subsequently, the developmental influences captured by *D* were removed from the data set of heat-stressed nematodes, allowing for the inference of the *heat-stress axis H. H* describes temporal expression patterns induced by heat stress, while disregarding heat-stress-independent temporal transcriptional patterns. Hence, by combining the data of perturbed and unperturbed populations, we were able to disentangle the effects of development and heat stress in time.

**Figure 1.**
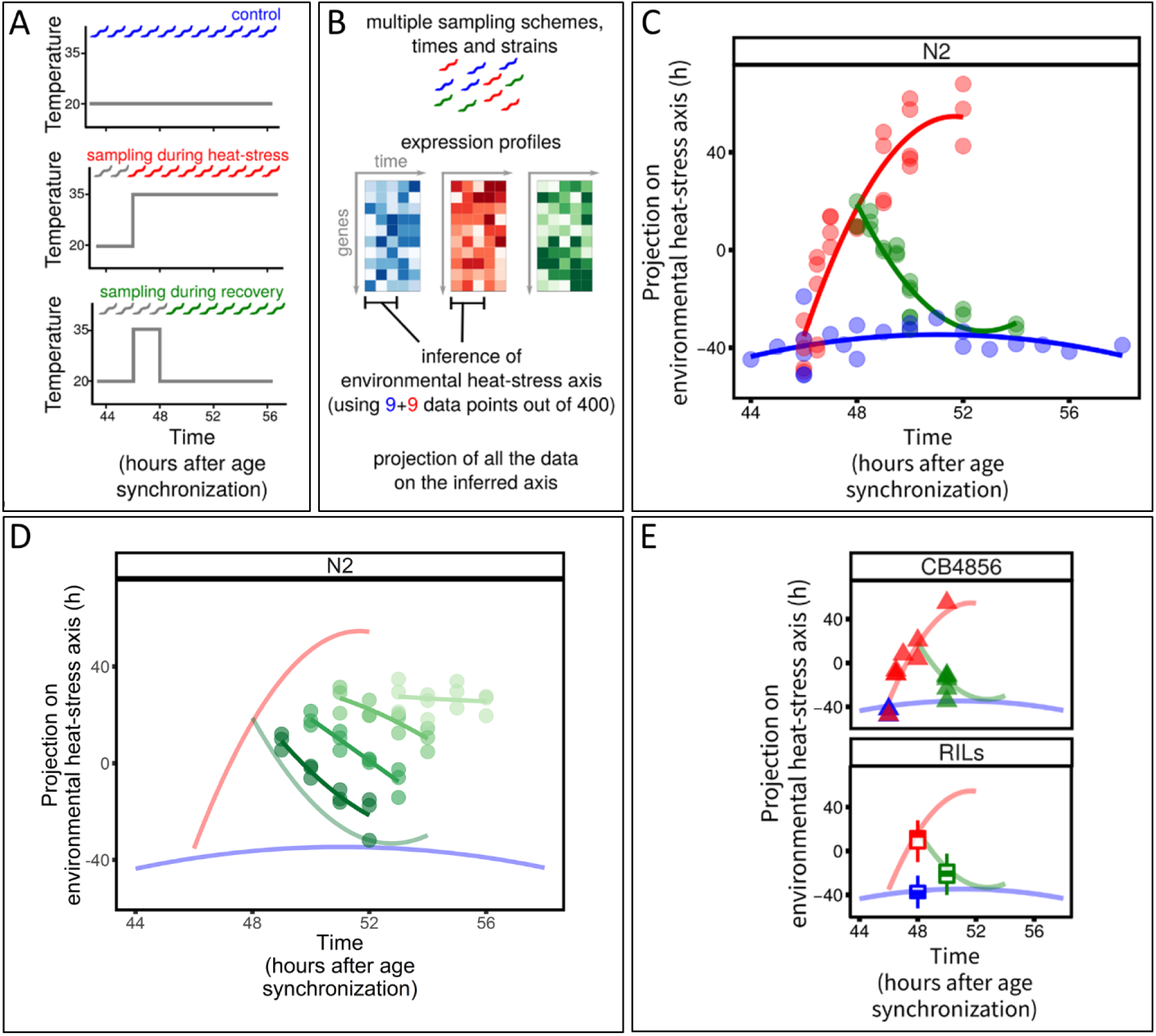
Experimental setup and expression dynamics during heat-stress perturbations. **A,**Experimental design of the three main treatments: control (blue; 20°C throughout development), heat stress (red; populations shifted to 35°C after 46 hours of development at 20°C), and recovery (green; at 20°C after heat stress). **B**, A subset of samples from heat-stress and control treatments were used for the inference of the heat-stress axis, *H*, describing the gene expression dynamics during heat stress. **C**, Projection of the data on this axis describes the dynamics of the response to heat stress. Notably, this is true also for the recovery data that was not used to infer axis *H*. **D**, Projection of transcriptome data of the recovery process after 2, 3, 4, and 6 hours of heat-stress shows a decrease in recovery dynamics. **E**, Axis *H* also describes the transcriptional heat-stress response for strains other than N2.

### Heat-stress axis *H* reflects exposure duration, as well as recovery from heat stress

By projecting gene-expression profiles on the heat-stress axis *H*, each sample can be associated with a value *h.* While only 18 samples were used to infer the axis, all 71 samples from all three time-series align along the axis according to treatment and exposure duration (Fig. 1C and Supplementary Figure S7), showing that *h* is a quantitative measure of the transcriptional stress response. The value of *h* increased with increasing heat-stress duration (Fig. 1C, red) until *h* started to saturate after long exposure (>4 hours). The unperturbed worms had a constant value of *h* (Fig. 1C, blue), showing that we were able to successfully remove the signal caused by developmental differences on gene expression. Strikingly, even though samples collected during recovery were not used to determine the axis *H*, the gene expression during recovery from a 2-hour heat stress was also well-explained with samples returning to the level of *h* typical of unperturbed worms within about four hours (Fig. 1C, green). We concluded that *h* quantitatively reflects exposure duration, as well as the time elapsed since the end of exposure. Note that although samples returned to the pre-stress treatment level of *h* after recovery, this does not imply that recovered *C. elegans* populations are transcriptionally indistinguishable from unperturbed ones (see SI section S2 and figure S7b). Therefore, recovery was defined and measured here by the ability to depart from stress response dynamics.

So far, the results have shown that the transcriptional recovery process after a mild stress can be followed over time using the heat-shock axis *H*. To exclude the possibility that *H* only captured time since the end of the heat stress without biological meaning towards phenotypic recovery or resilience, we expanded the dataset to include four additional time-series tracking the transcriptome recovery for 4 hours following four different heat-stress intensities (2, 3, 4, and 6 hours at 35°C). The long-term effect of these stress intensities on survival, reproduction, and mobility have been shown to range from mild after short (2 hour) exposure to 100% mortality within 24 hours after 6 hours at 35°C [8]. Fig. 1D shows that mildly stressed population transcriptionally returned to pre-stress levels of *h* during the observed recovery period, while increasing stress duration led to a slowing down of the transcriptional recovery process, and severely stressed populations remained at a constant high value of *h.* Therefore, *H* can distinct between the progress of the recovering transcriptome and a non-recovering transcriptome.

### Heat-stress axis *H* retains essential features of the biology of the heat-stress response

Having shown that the axis *H* recapitulates the transcriptional state during and after exposure to heat stress, we investigated the biological properties of the axis *H*. To this end, we performed an enrichment analysis to determine which groups of genes contributed the most to the axis *H* (Fig. 2). Consistent with expectations, genes encoding for stress response proteins (in particular heat-shock proteins *hsp*) and nucleosomes (in particular histones *his*) were upregulated. These gene classes have previously been shown to be highly activated by heat stress [7–9, 14]. Similarly, the value of *h* is negatively correlated with some genes involved in cell metabolism such as ATPase transmembrane proteins. This analysis shows that the axis *H* retains essential features of the biology of the heat-stress response, while summarizing these complex biological dynamics into a single quantitative parameter.

**Figure 2.**
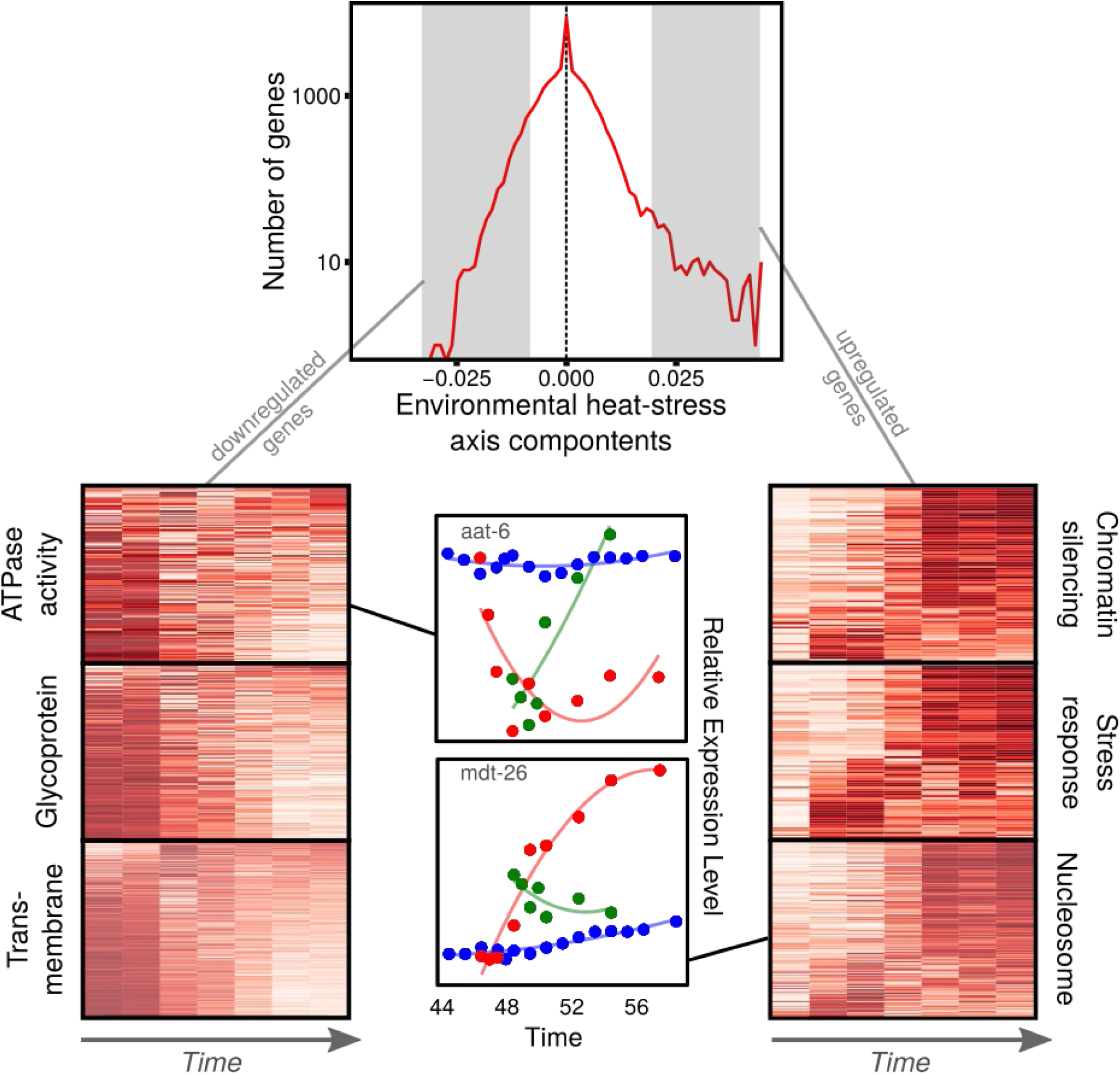
Single gene contribution to heat-stress axis, *H*. Distribution of the entries of the heat-stress axis (top of the figure). The distribution is not symmetric, which means that most genes (relatively to their unperturbed level of expression) are upregulated during heat-stress response. Enrichment results including two example of genes with negative and positive components (red corresponds to expression levels measured during heat stress, green corresponds to recovery, and blue to development).

### Heat-stress axis *H* reflects the average heat-stress response across multiple genotypes

Next, we tested whether heat-stress axis *H* can also reflect the change in gene expression for different genotypes. We used expression profiles of the strain CB4856 (Hawaii), which is genetically distinct from N2, as well as 54 recombinant inbred lines (RILs) [9], which are genetic mosaics derived from a cross between CB4856 and N2 [15, 16]. We found that the heat-stress axis *H* successfully recapitulates the average dynamic transcriptional response and resilience of this genetically diverse set of lines (Fig. 1E and SI section S3). The robustness of the pattern across genotypes reflects the high degree of conservation in transcriptional resilience. It should be noted that the RILs were not used here for the genetic mapping of traits [17], but rather as a genotypic library.

### Variation in stress resilience across genotypes is captured in a genetic heat-stress axis (*GH)*

We have shown that the heat-stress axis *H*, inferred using solely the isogenic strain N2, describes the *average* conserved stress response of a library of highly divergent genotypes. On the other hand, there is large natural variation in long-term effects of heat-stress exposure across genotypes, for example marked by differences in the stressed transcriptome [9], survival rates [3, 18], and reproductive rates [3]. Considering that variation is genotype dependent, it implies a difference in transcriptional resilience during and/or after stress. Next, we ask if a single axis could also capture the natural variation in heat-stress response across genotypes. Since genotypes differ in more traits than their transcriptional response to stress, such as developmental timing and size, we needed to isolate stress-induced variation in expression levels from other intrinsic differences in the transcriptome between genotypes. For this purpose, we used gene expression data of RILs collected before and after two hours of heat stress [9]. Analogous to our approach above in inferring the heat-stress axis *H* for N2 by removing developmental differences, we corrected the heat-stress response of the RILs for their intrinsic gene expression differences in unperturbed conditions (see SI section S3). We inferred a *genetic heat-stress axis* (*GH*) that isolates and describes the variability across strains in their stress response.

The strength of relationship between the genetic axis *GH* and the environmental heat-stress axis *H* measures the proportion of the variation of heat-stress response across RILs that is due to timing differences. We found a positive correlation between the two axes (Spearman rho = 0.36, p = 0.01) implying that different strains respond as if they were exposed to the heat stress for different durations. This was confirmed by analysing a second set of heat-stressed gene expression profiles from a separate alternative panel of inbred lines [19, 20](Introgression Lines, ILs; Spearman rho = 0.44, p = 8·10^−4^) (Fig. 3). These results show that the genetic differences also lead to difference in the timing or magnitude of the transcriptional response. The presence of a correlation between the axes *H* and *GH* also implies that the value of *gh* (the projection of the gene expression profile on the axis *GH*) recapitulates the relative strength of the heat-stress response: the higher the value of *gh*, the farther away is the gene expression profile from the unperturbed state or, in other words, the lower its transcriptional resilience to heat-stress.

**Figure 3.**
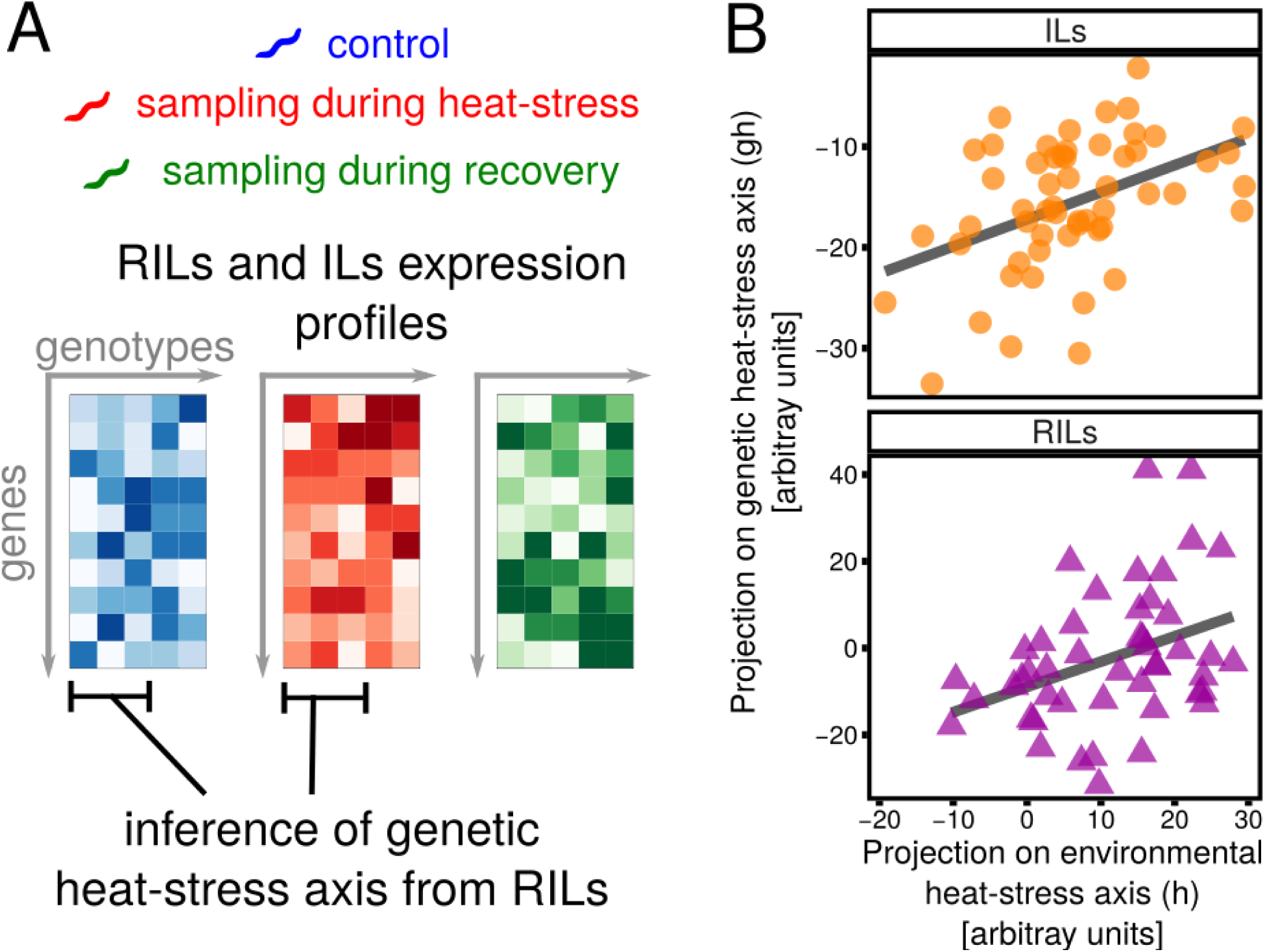
Derivation of the genetic heat-stress axis, *GH*, and relation with environmental heat-stress axis, *H.* **A**, For each RIL and IL, we measured gene expression in absence of perturbation, after two hours exposure to heat stress and during recovery (after two hours of the end of a two-hour heat stress). Using only RIL data, we obtain the genetic heat-stress axis (*GH*), describing the difference between RILs in heat-stress response (discounting their differences in gene expression in the unperturbed case). **B**, Correlation between genetic heat-stress axis and the environmental heat-stress axis shown for heat-stress samples of RILs and ILs.

### Transcriptional resilience on a short timescale is predictive of the variability in thermotolerance on a longer timescale

Heat stress affects gene expression dynamics and resilience in the short term in a predictable way, which is recapitulated by axes *H* and *GH*. On the other hand, in the long run, heat stress also affects developmental speed, aging, behaviour, and vitality - for instance by drastically reducing lifespan. We set out to explore how the variability in gene expression dynamics following heat stress on a short timescale is predictive of variability in thermotolerance measured on a longer timescale. Thermotolerance in *C. elegans* can be recorded by its survival rates. Therefore, in a parallel experiment, we collected lifespan data of over 200 different RILs and ILs with and without exposure to heat stress. While two hours at 35°C are sufficient to induce a strong transcriptional response, previous experiments have shown that overall lifespan is not necessarily shortened at this intensity [8]. Therefore, we increased the exposure duration to four hours at 35°C for lifespan measurements as this duration is known to affect lifespan [3, 8], allowing us to make a better estimate of difference in thermotolerance across genotypes As expected, both RILs and ILs show high variability in their lifespan after heat stress and in control conditions. The average lifespan following a heat stress is always lower than what was found for the same strain when unperturbed (Fig. 4 and Figure S14).

**Figure 4.**
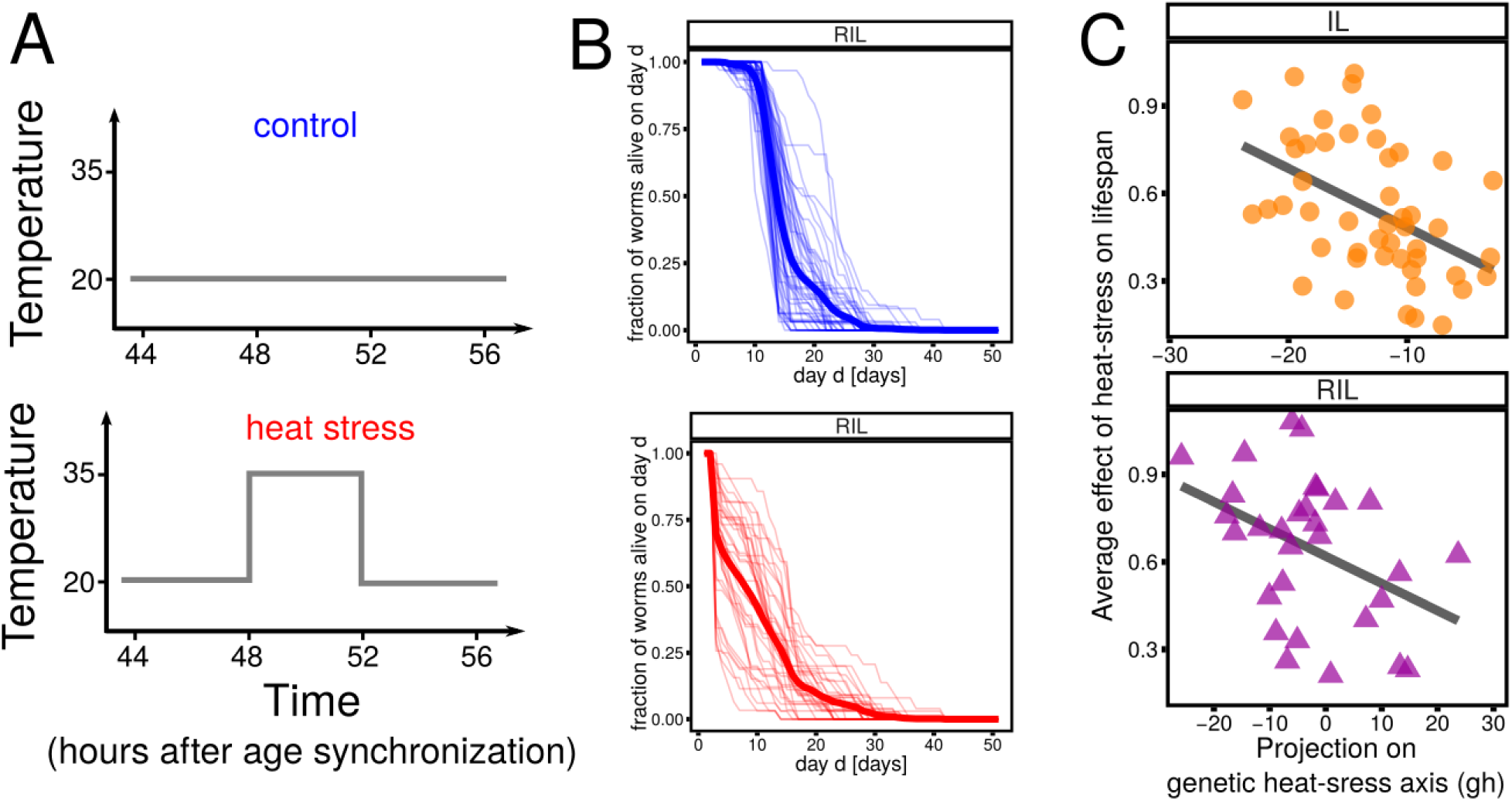
Effect of heat stress on lifespan and correlation with gene expression recovery. **A**, Experimental set-up used to collect lifespan data of 40 RILs and 54 ILs. **B**, Comparison of the cumulative lifespan distribution of unperturbed (blue) and perturbed (red) RILs. Each thin line corresponds to a RIL (an average of 31 animals were scored per genotype and treatment), while the thick line is the average across RILs. **C**, Effect of heat stress on lifespan (measured as the ratio of the average lifespan after perturbation and without perturbation) correlates with the projection of recovery data on the genetic heat-stress axis for RILs and ILs. Strains recovering faster from heat stress experience a weaker effect on their lifespan.

Next, we compared the effect of heat stress on the lifespan of different RILs with the difference in transcriptional resilience, measured by projecting the recovery data of the RILs on the genetic heat-stress axis *GH*. Figure 4 shows that the ability of different strains to recover from heat stress is predictive of thermotolerance (Spearman rho = −0.41, p = 0.02; Fig 4). In order to test the robustness of this result, we also performed the same analysis on ILs, which are genetically mostly derived from one strain (N2) and were not used to infer the axis. In this case, we also found a significant correlation (Spearman rho = −0.46, p = 10^−3^), implying that the connection between the ability to recover and lifespan was robust across different inbred line panels. The projection of the heat-stress data onto *GH* (which is related to the speed at which worms react to heat stress) was not robustly correlated with lifespan (see SI section S5), showing that resilience measured based on recovery data was more directly linked to tolerance.

## Conclusions

This study sheds light on how organisms recover from environmental stress perturbations, by means of a systemic *modus operandi* based on using genome-wide gene expression profiles. We conclude that a relatively simple axis can measure stress resilience of a dynamic transcriptome in a single quantitative variable and describes the capacity of an organism to recover from heat stress. Our findings show that natural variation in transcriptome resilience after mild stress exposure is predictive of thermotolerance across a diverse set of genotypes in *C. elegans*. The results imply that thermotolerance is an intrinsic trait that largely pre-determines long-term effects of heat-stress exposure. Operationalizing the concept of resilience in higher organisms, like mammals, has been difficult because it includes a range of many different phenotypic traits [21]. Our approach represents a novel way in understanding resilience in a living system, and we show how the inherent complexity of stress recovery can be exploited to predict the chance of survival. We anticipate that our finding will accelerate progress in the study of resilience of complex living systems, opening up new avenues of research in stress, aging, and disease across other species.

## Methods

### Strains and Maintenance

The wild-type *C. elegans* strains N2 (Bristol) and CB4856 (Hawaii) were used in this study, as well as 54 CB4856 × N2 recombinant inbred lines (RILs; each line is a genetic mosaic with contributions of the two parental strains [9, 15, 16], and 47 CB4856 × N2 introgression lines (ILs; one individual locus of the CB4856 genome introgressed into an otherwise N2 genetic background [19]). Strains were maintained under standard culturing conditions on (9 cm diameter) Petri dishes with Nematode Growth Medium (NGM) containing *Escherichia coli* OP50 as food source [22]. To prepare populations for the start of an experiment, maintenance populations were chunked to fresh 9-cm NGM plates with food and kept at 20°C for exactly one week to induce starvation. This was done to assure that all populations received the same treatment before the experiment.

### Lifespan under heat stress and control conditions

Starved populations were transferred to fresh NGM dishes seeded with *E. coli* OP50 by chunking, and grown at 16°C or 20°C (depending on the desired growth rate) for 3-4 days to obtain proliferating populations. The populations were age synchronized by hypochlorite treatment according to standard protocols [22] and grown at 20°C until the 4th larval stage was reached. At 47 hours post age-synchronization, the larvae were collected from the plates with M9 buffer, and 30-40 individuals were transferred to new NGM dishes containing 5-fluorodeoxyuridine [23]. At 48 hours post age-synchronization, the heat-stress treated group was exposed to 35°C for 4 hours. Control and post-heat-stress conditions were set to 20°C. Survival was scored every day by checking the response to touch with a picking needle. For each genotype, an average number of 31 animals were scored for each treatment.

For each genotype and treatment the survival curves are reported in Figure 4B. For each genotype and treatment, we computed the average lifespan. The average effect of heat stress on lifespan (reported in Fig. 4C) is the ratio between the average lifespan after heat stress and the average lifespan in the control.

### Transcriptome profile of heat stress, recovery and developmental state

#### Data retrieval

Sub-sets of the transcriptomic profiles used in this study have previously been described. These sub-sets include the developmental time series previously described in Snoek *et al.* (2014) [12], retrieved from Array Express https://www.ebi.ac.uk/arrayexpress/ under accession number E-MTAB-7019, which includes 22 transcription profiles of N2 populations sampled in hourly intervals between 44-58 hours post synchronization. The heat-stress time series was described in Jovic *et al.* (2017) [8], which includes 29 transcription profiles established after an exposure to 35°C for 0, 0.5, 1, 2, 3, 4, 6, 8, or 12 hours starting at 46 hours after age-synchronization (retrieved from ArrayExpress under accession number E-MTAB-5753). Transcription profiles of the RILs and ILs including the parental lines (N2 and CB4856) in control conditions (sampled 48 hours post age-synchronization), after a 2-hour heat stress starting at 46 hours post age-synchronization, and after a subsequent 2-hours recovery period were first presented by Snoek *et al.* ([9]; data retrieved from ArrayExpress, accession number E-MTAB-5779) and Sterken *et al.*, ([20]; ArrayExpress, accession number E-MTAB-7424), respectively. A detailed overview of data sets and the according publications can be found in Supplementary table 1.

#### Stress treatment and sampling for transcriptome analysis

The above described transcriptome data-set was extended with a heat-stress time series for CB4856, and a recovery time series for N2 using protocols adapted from Snoek *et al.* (2014, 2017) [9, 12] and Jovic *et al.* (2017) [8]. Starved populations were transferred onto fresh NGM dishes seeded with *E. coli* OP50 by chunking and grown at 16°C or 20°C for 3-4 days depending on the desired growth rate to obtain gravid hermaphrodites. Age-synchronized populations were obtained by hypochlorite treatment according to standard protocols [22] and maintained at 20°C until the begin of the heat-stress exposure of 35°C starting at 46 hours post age synchronization. For the CB4856 heat-stress time series, samples were taken after 0, 0.5, 1, 2, 4, and 6 hours at 35°C by rinsing the populations of the NGM plates with M9 buffer. For the N2 recovery time series, samples were transferred to 20°C after 2, 3, 4, or 6 hours at 35°C. After 2 hours of heat-stress, samples were taken 0, 0.5, 1, 1.5, 2, 4, and 6 hours into the recovery period; For 3, 4, or 6 hours of heat stress, samples were taken in an hourly interval up to 4 hours post-exposure. All samples were immediately flash frozen in liquid nitrogen at the time of collection, and stored at −80°C until further use.

#### RNA isolation

mRNA was isolated from frozen samples using the Maxwell® 16 AS2000 instrument with a Maxwell® 16 LEV simplyRNA Tissue Kit (both Promega Corporation, Madison, WI, USA). The mRNA was isolated according to protocol with a modified lysis step as described in Snoek *et al.* 2014, and Jovic *et al.* 2017. 200 μl homogenization buffer, 200 μl lysis buffer and 10 μl of a 20 mg/ml stock solution of proteinase K were added to each sample. The samples were then incubated for 10 minutes at 65°C and 1000 rpm in a Thermomixer (Eppendorf, Hamburg, Germany) before cooling on ice for 1 minute. At this point, the samples were pipetted into the cartridges resuming with the standard protocol.

#### Sample preparation and scanning

For cDNA synthesis, labelling and the hybridization reaction, the ‘Two-Color Microarray-Based Gene Expression Analysis; Low Input Quick Amp Labeling’ - protocol, version 6.0 from Agilent (Agilent Technologies, Santa Clara, CA, USA) was followed, starting at step 5. The Agilent *C. elegans* (V2) Gene Expression Microarray 4X44K slides were used in combination with an Agilent High Resolution C Scanner using the recommended settings. Data was extracted with the Agilent Feature Extraction Software (version 10.7.1.1) following the manufacturers’ guidelines.

#### Data normalization

Microarray data were normalized using a within array normalization using a standard function of the R package limma (using “loess” method) [24].

The code needed to replicate all the results presented here can be found at https://github.com/jacopogrilli/resiliencevitality.git

## Availability of data and materials

Newly produced raw and processed transcriptome datasets supporting the conclusions of this article are available in the ArrayExpress database at EMBL-EBI (www.ebi.ac.uk/arrayexpress; [25]) under the accession number E-MTAB-7007 and E-MTAB-7948.

## Acknowledgements

We thank Miriam Rodriguez, Roel P. J. Bevers, Rita Volkers, Rosanne Bartels, Yvonne Laven for the assistance with collecting data. This research was financed by the Human Frontiers Science Program (grant RGP0028-2014). We thank Ben Lehner, Riekelt Houtkooper, Marco Cosentino Lagomarsino, Zachary Miller, and Wim van der Putten for helpful discussions about the manuscript.

## Author Contributions

KJ, JG, SA and JEK wrote the manuscript. MGS and BLS commented on the manuscript. KJ, MGS and JAGR performed and supervised the experiments. KJ, MGS and BLS assembled, annotated and curated the microarray data. JG performed the analysis and wrote the code.

