## Supplemental Information 1 for "Transcriptome dynamics predict thermotolerance in *Caenorhabditis elegans*"

### Supplementary Information

**Contents**


**S1. Singular Value Decomposition and Principal Component Analysis** 2

**S2. Principal axis for N2** 3

A. Derivation of the developmental axis D 3

B. Derivation of the heat-stress axis H 5

C. Projection of the recovery data onto H 8

**S3. RILs data** 8

A. Projection of RILs data on H 8

B. Derivation of the genetic heat-stress axis, GH 11

**S4. Lifespan data** 12

**S5. Heat-stress response and lifespan** 13

**References** 16

#### S1. SINGULAR VALUE DECOMPOSITION AND PRINCIPAL COMPONENT ANALYSIS

In this section, we describe how we obtained the principal axis from the data. After the canonical corrections of the microarray data (see Methods), we can arrange our data in a matrix $\mathbf{X}$, with $G$ rows and $S$ columns, where $G$ is the number of spots in the microarray ($G = 45220$) and $S$ is the number of samples. The element $X_{sg}$ represents the logarithm of the intensity ("expression level" henceforth) obtained from the micro-array data after the procedures described in the Methods.

Since the variability in both the average and the variance of intensities across the samples might not be due to biological differences, we rescale the data such that the average expression level is equal to 0, and the variance is equal to 1. The transformation is

$$\tilde{X}_{sg} = \frac{X_{sg} - (1/G)\sum_g X_{sg}}{\sqrt{(1/G)\sum_g X_{sg}^2 - \left((1/G)\sum_g X_{sg}\right)^2}} \; , \tag{1}$$

such that

$$\frac{1}{G}\sum_g \tilde{X}_{sg} = 0 \; , \tag{2}$$

and

$$\frac{1}{G}\sum_g \tilde{X}_{sg}^2 = 1 \; . \tag{3}$$

In full generality, the elements of the matrix $\tilde{\mathbf{X}}$ can be written as

$$\tilde{X}_{sg} = \sum_{\alpha=1}^{S} \lambda_\alpha w_s^\alpha v_g^\alpha \; , \tag{4}$$

where

$$\sum_{s=1}^{S} w_s^\alpha w_s^\beta = \sum_{g=1}^{G} v_g^\alpha v_g^\beta = \delta_{\alpha\beta} \; , \tag{5}$$

and $\delta_{\alpha\beta}$ is the Kroneker delta (equal to one if $\alpha = \beta$, and zero otherwise). The factorization of the matrix $\tilde{\mathbf{X}}$ shown in equation 4 is called the Singular Value Decomposition (SVD) of the matrix $\tilde{\mathbf{X}}$. The vectors $\mathbf{w}^\alpha$ and $\mathbf{v}^\alpha$ are the singular vector of the matrix $\tilde{\mathbf{X}}$—specifically, one can define the $S \times S$ matrix $\tilde{\mathbf{X}}^T\tilde{\mathbf{X}}$ which has eigenvalues $\lambda_\alpha^2$ and eigenvectors $\mathbf{w}^\alpha$ (note that this matrix is positive semi-definite, as it can be seen as a covariance matrix). On the other hand, the $G \times G$ matrix $\tilde{\mathbf{X}}\tilde{\mathbf{X}}^T$ has $S$ eigenvalues equal to $\lambda_\alpha^2$ with eigenvectors $\mathbf{v}^\alpha$, while the other $G - S$ eigenvalues are all equal to zero.

The entries of the matrix $\tilde{\mathbf{X}}^T\tilde{\mathbf{X}}$ are defined as

$$(\tilde{\mathbf{X}}^T\tilde{\mathbf{X}})_{ss'} = \sum_g \tilde{X}_{sg}\tilde{X}_{s'g} \; . \tag{6}$$

Because of the relation in equation 3, the diagonal elements are all equal to one, while the off diagonal coefficients represent the correlations between the expression pattern of two samples. The diagonalization of this matrix and $\tilde{\mathbf{X}}\tilde{\mathbf{X}}^T$ allows to identify independent combinations of genes that explain most of the variation observed in the data.

Principal Component Analysis employs singular value decomposition (i.e., the factorization shown in equation 4) to identify the components that explain most of the variation in the data. Since all the values $\lambda_\alpha$ are positive, one can always ordered them to have $\lambda_1 > \lambda_2 > \cdots > \lambda_S \geq 0$. The first principal component (often called PC1) is then simply $w_s^1$, the second one $w_s^2$ and so on. The fraction of variance explained by the $k$th component can be simply expressed in terms of the eigenvalues

$$\frac{\lambda_k^2}{\sum_{\alpha=1}^{S} \lambda_\alpha^2} \ . \tag{7}$$

The vectors $\mathbf{v}^\alpha$ parallel the information given by the vectors $\mathbf{w}^\alpha$. Two samples , $s_1$ and $s_2$, have similar expression patterns if the entries $s_1$ and $s_2$ of the first principal components $\mathbf{w}^1$, $\mathbf{w}^2$, ... have similar values. In the same way, two genes $g_1$ and $g_2$ have similar expressions *across* samples, in the corresponding entries of $\mathbf{v}^1$, $\mathbf{v}^2$, ... are close. It is important to remember that, if the vector $\mathbf{v}^\alpha$ is known, the corresponding vector $\mathbf{w}^\alpha$ can be obtained from

$$\sum_g \tilde{X}_{sg} v_g^\alpha = \lambda_\alpha w_s^\alpha \ . \tag{8}$$

#### S2. PRINCIPAL AXIS FOR N2

In this section, we explain how we obtain the developmental axis, D, and the heat-stress response axis, H, for N2.

##### A. Derivation of the developmental axis D

The developmental data of the N2 strain were obtained in two batches, *set*11 (12 data points) and *set*13 (8 data points), see methods. We used only the data from the batch *set*11 to infer the principal axis $\mathbf{v}^\alpha$. Once the axis is known, one can obtain the corresponding component by projecting the data over these vectors (see equation 7). In this section, we show that—using only the gene expression levels of one batch (without using a-priori information on the developmental stage of individual samples)—we are able to describe the change in expression due to development of all the other samples in the full dataset.

As explained in section S1, we consider the data matrix $\tilde{\mathbf{X}}$, whose entry $\tilde{X}_{sg}$ represents the normalized logarithm of the expression level of gene $g$ in sample $s$.

Figure S1 shows that the distribution of gene expression levels across different samples is bimodal in both batches. For all samples, the minimum of the distribution between the two maxima occurs at an expression level around 4.5. Since genes that have low expression are noisier, we removed, for the purposes of the derivation of the axis, all

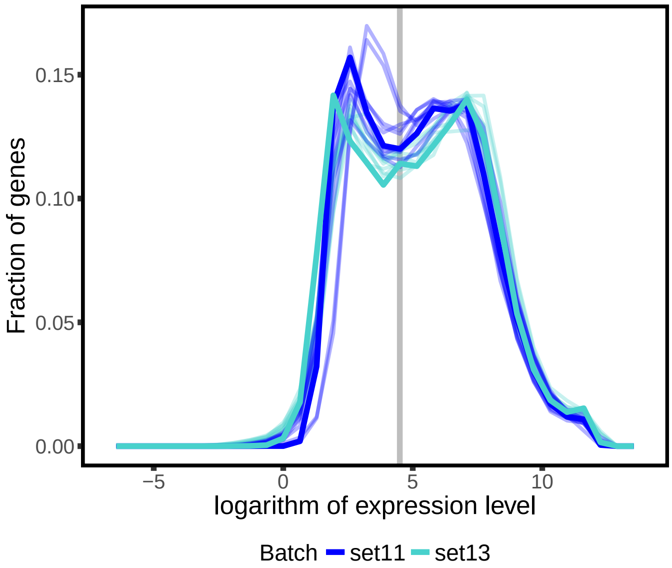


Supplementary Figure S1: Distribution of the logarithm of gene expression levels of developmental data. Each curve in the background corresponds to a sample, while the two solid curves mark the average over the samples of a given batch (dark blue for *set*11 and light blue for *set*13). While different samples correspond to different stages in development, the distribution is quite robust and is always bimodal. The gray line correspond to a value of the logarithm of the expression level equal to 4.5, which is the value we use as a threshold between highly- and lowly-expressed genes.

the genes whose average expression across the samples of batch *set*13 was lower than 4.5. In practice, we set the corresponding entries of $v_g^\alpha$ to zero.

Using the genes with average log-expression larger than 4.5, we removed mean and variance obtaining the data matrix $\tilde{\mathbf{X}}$ as described in section S1. Using only the samples from *set*13, we performed the singular value decomposition obtaining the principal axis $\mathbf{v}^\alpha$. In order to obtain the $k$-th principal component of any sample from any batch, we need to project the gene expression data on the axis $\mathbf{v}^k$. In other words, if $\tilde{X}_{sg}$ is the logarithm of the expression of gene $g$ in sample $s$, the projection of sample $s$ on the $k$-th axis is given by $\sum_g \tilde{X}_{sg} v_g^k$.

The first principal axis $\mathbf{v}^1$ is strictly related to the heterogeneity of expression levels among genes. Figure S2 shows that the entry of the principal vector is correlated with the average expression of genes. In particular, the relation is perfectly linear for batch *set*13, which is the one used to infer the axis. The first principal axis measures therefore the average expression and not the difference between samples and their change in time. Figure S2 shows that the second principal axis, on the other hand is not correlated with the average expression, implying that this vector captures other structural properties of the data.

Figure S3 shows that the projection of the data on the second principal axis has a clear trend with time (*i.e.* developmental age measured in hours after age-synchronization, see methods for details). As such, the second principal axis is related to the developmental stage of age-synchronized N2 populations. It should be noted that the develop-

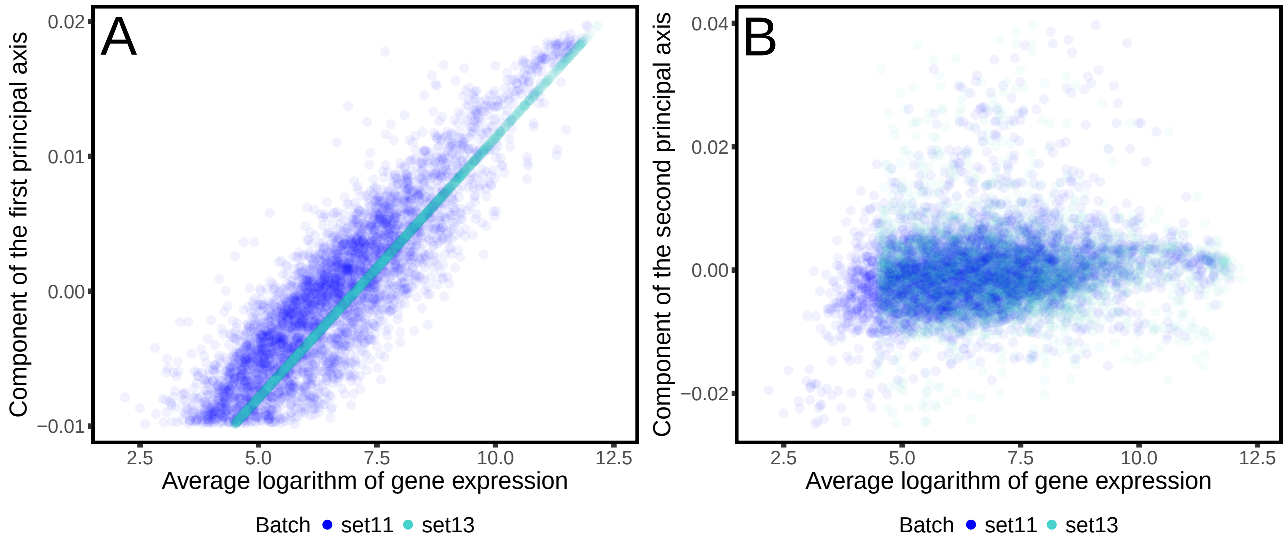


Supplementary Figure S2: Correlation between the principal axes and gene expression. The two panels represent the component of the first (panel A) and second (panel B) principal axis vs. the corresponding average expression level. The two axes were inferred using only the data from batch *set*13 sampled during unperturbed development. Panel A shows that the component of the first axis perfectly correlates with the average expression level of the corresponding gene for *set*13. This observation implies that the first principal axis only summarizes the average expression level of different genes and does not include any information on the dynamics of gene expression. Panel B shows that instead the second principal axis components are not correlated with average gene expression.

mental axis, D, is also relating to the developmental timing of other samples within other batches, and also for other genotypes (see Figure S3), even though it was obtained using only the samples of batch *set*13.

##### B. Derivation of the heat-stress axis H

To produce the time-series profiling the expression levels during the response to heat-stress, N2 populations were exposed to 35°C for different durations (see methods for more details). Data were collected in 7 batches. In order to obtain the heat-stress axis, we applied the same procedure as described in section S2 A.

First, we normalized the data, obtaining the matrix $\tilde{\mathbf{X}}$. Then, we derived the principal axis only considering the batch *set*1 (10 data points out of a total 39 heat-stressed N2 samples) and only the genes which had a log expression level averaged over the samples within batch *set*1 higher than 4.5.

The components of the first principal axis determined with this procedure are again correlated with the average gene expression and do not capture the information on the effect of different heat-stress perturbations (see Figure S4). As in the case of the developmental axis, we expect the second principal axis to be related to the time elapsed since the beginning of the perturbation. While the worms were exposed to heat-stress starting at a given age (*i.e.*, 46h

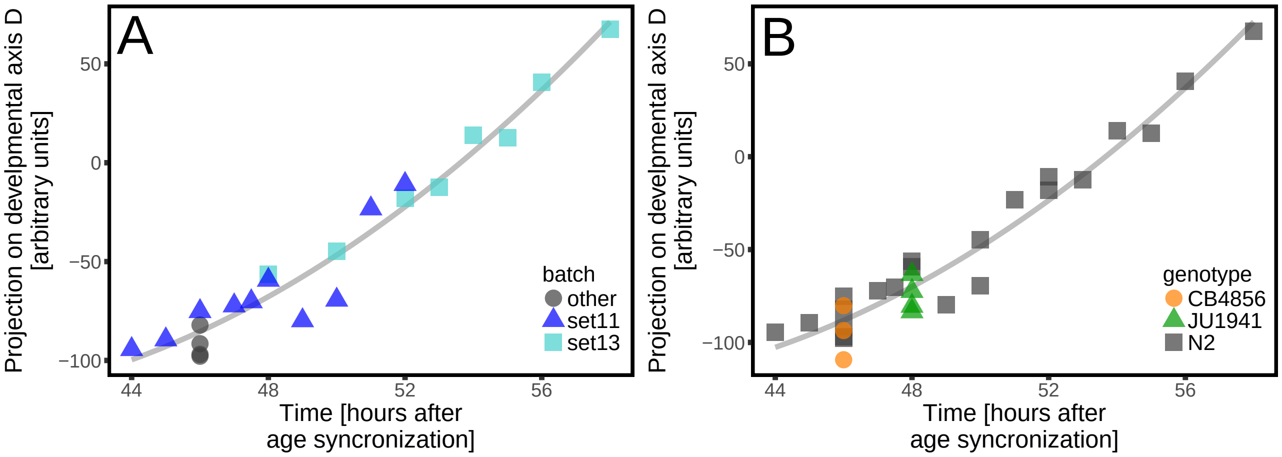


Supplementary Figure S3: Projection on the second principal axis (developmental axis, D) vs. time. The projection on the developmental axis shows a clear dependence on time, suggesting that the complex multi-dimensional dynamics of gene expression during development can efficiently be reduced to a single variable. Panel A shows that the axis, which was inferred using only the data from one batch (*set*13), also describes the dynamics of the other batch (*set*11). Panel B compares the projection of the N2 data (grey squares) with the projection of the strains CB4856 (orange circles) and JU1941 (green triangles), showing that the developmental axis also describes the expression dynamics of other strains.

post age-synchronization), development partially continues during heat-stress [Snoek *et al.*, 2017]. Therefore different exposure durations to heat-stress also correspond to different developmental ages. In other words, the observed effect on worms exposed to a heat-stress for different times is the combined effect of heat-stress response and development. In order to disentangle the effect of development from the effect of heat-stress, we considered the second principal axis found from *set*1 and we removed from it its projection on the developmental axis. The heat-stress axis, H, found in this way is orthogonal (and therefore independent) to the developmental axis, and captures only the effect of different heat-stress exposures.

Figure S5 shows the projection on the heat-stress axis of individual populations exposed for different lengths of time—demonstrating how the axis is related to the heat-stress exposure duration. It should be noted that the axis was obtained using only a small sub-set of data points in *set*1, but it well describes the heat-stress response of all data points within the other sets. Moreover, the axis also describes the effect of heat-stress on the gene expression of other strains than N2. Therefore, the heat-stress axis H provides a robust and quantitative measure of the heat-stress response across different genotypes.

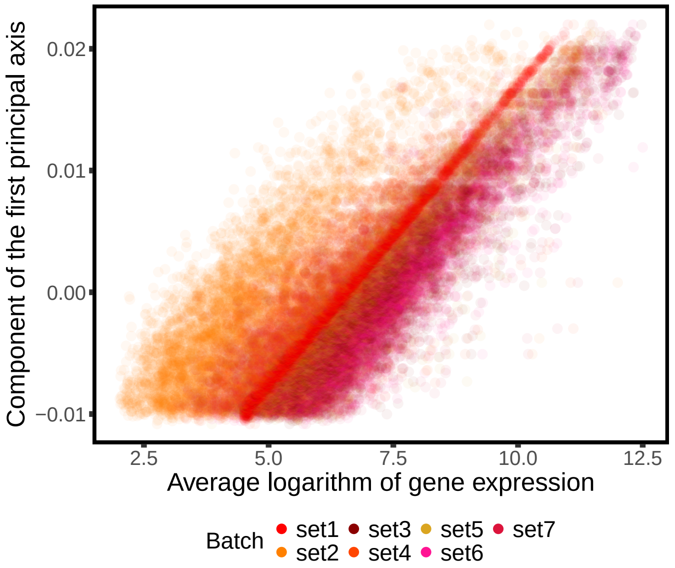


Supplementary Figure S4: Correlation between first principal axis and gene expression. The axis was inferred using only the heat-stress data from batch *set*1. The component of the first axis correlates perfectly with the average expression level of the corresponding gene for *set*1, and it correlates strongly with average expression for the other batches. As such, the first principal axis only summarizes the average expression of different genes and does not include any information on the dynamics of gene expression.

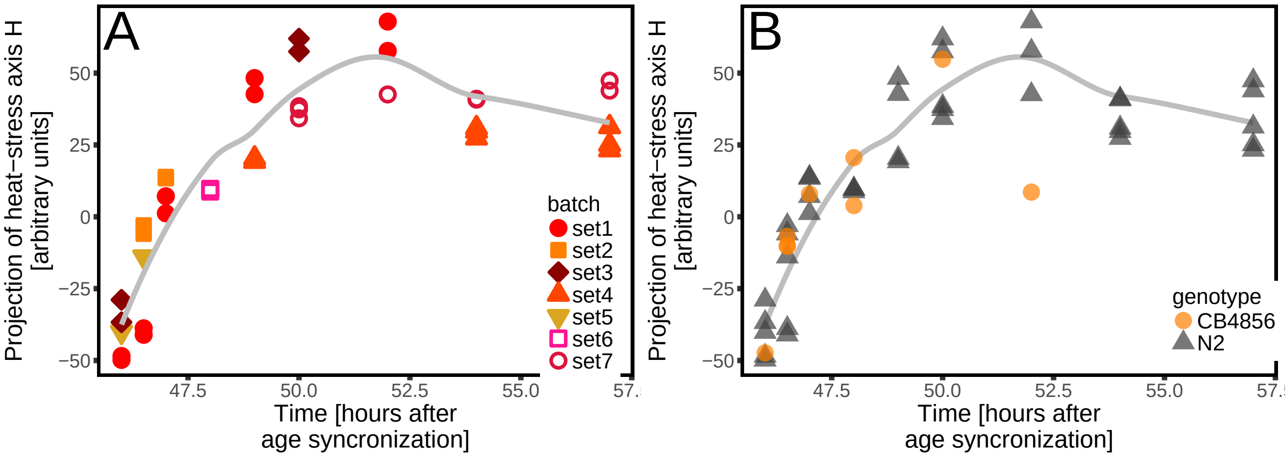


Supplementary Figure S5: Projection of the heat-stress data on the second principal axis (heat-stress axis, H) vs. time (note that heat-stress was started at 46 hours). The projection on the heat-stress axis shows a clear dependence on time, suggesting that the complex multi-dimensional dynamics of gene expression during heat-stress can be efficiently reduced to a single variable. Panel A shows that the axis inferred only by using the data from one batch (*i.e.*, *set*1) also describes the dynamics of other batches. Panel B compare the projection of the N2 data (gray triangles) with the projection of the strain CB4856 (orange circles), showing that the heat-stress axis H also recapitulates the heat-stress response of other strains.

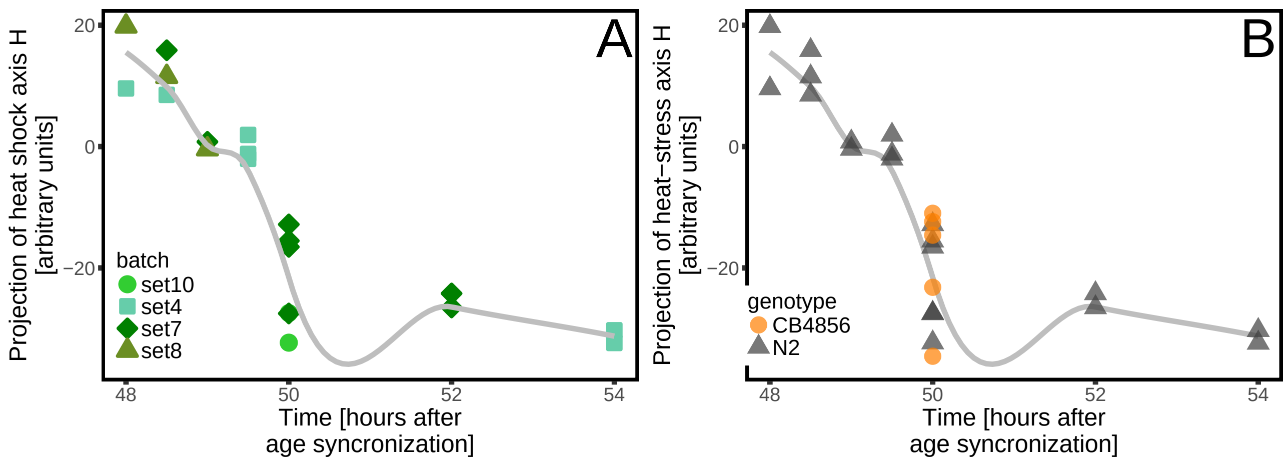


Supplementary Figure S6: Projection of the recovery data on the heat-stress axis vs time. The heat-stress axis was inferred by using only one batch (*i.e.*, *set*1), obtained by exposing the worms to a heat-stress for different times. Importantly, the axis also describes the recovery after heat-stress (panel A), implying that recovery follows a pattern similar to the response. Moreover, panel B shows that the axis also describes the recovery of the strain CB4856 (orange circles).

##### C. Projection of the recovery data onto H

To analyze the gene expression during recovery, N2 populations were exposed to two hours at 35°C degrees, after which they were put back to 20°C degrees. Samples were taken at different times during the recovery period (see methods for details). Figure S6 shows the projection of the recovery data on the heat-stress axis, showing that H also describes the recovery after perturbation. This is notable, given that we did not use the recovery data to infer the heat-stress axis H, nor to infer the developmental axis D.

Figure S7 shows the projection of the gene expression level of the parental lines on the heat-stress axis and developmental axis, summarizing which data points were used to infer the axes.

#### S3. RILS DATA

##### A. Projection of RILs data on H

To take a deeper look at genetic variation in the heat-stress response, we used gene expression profiles of a genetically diverse set of recombinant inbred lines (RILs) and introgression lines (ILs) in unperturbed development, after 2 hours heat-stress, and after 2 hours of recovery (see methods for more details). Figure S8 shows the projection of the RILs and ILs data on the heat-stress axis. RILs and the ILs samples have similar projections on H before, during and after the heat-stress. Moreover, the values of their projections is not particularly different from the values of the parental lines N2 and CB4856.

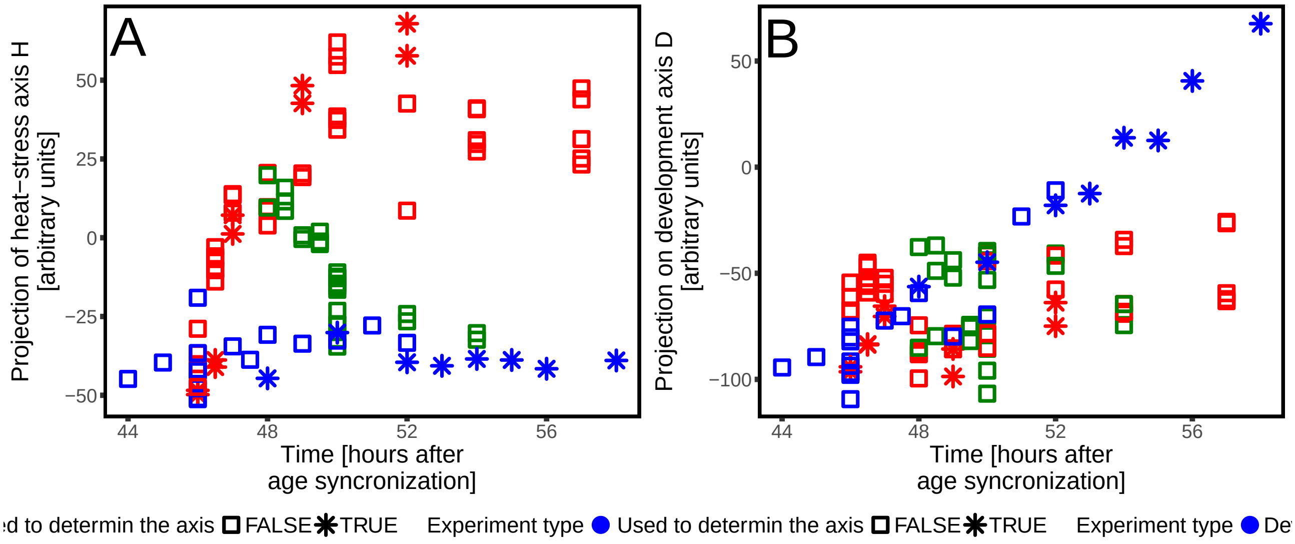


Supplementary Figure S7: Projection of unperturbed, perturbed and recovery data on the heat-stress axis H (panel A) and developmental axis D (panel B) vs time. Different colors correspond to different treatments (blue are unperturbed worms, red corresponds to the heat-stress data and green is recovery data). Only the data points marked with a star were used to determine the two axes, while the square symbols correspond to data that was not used in the inference process.

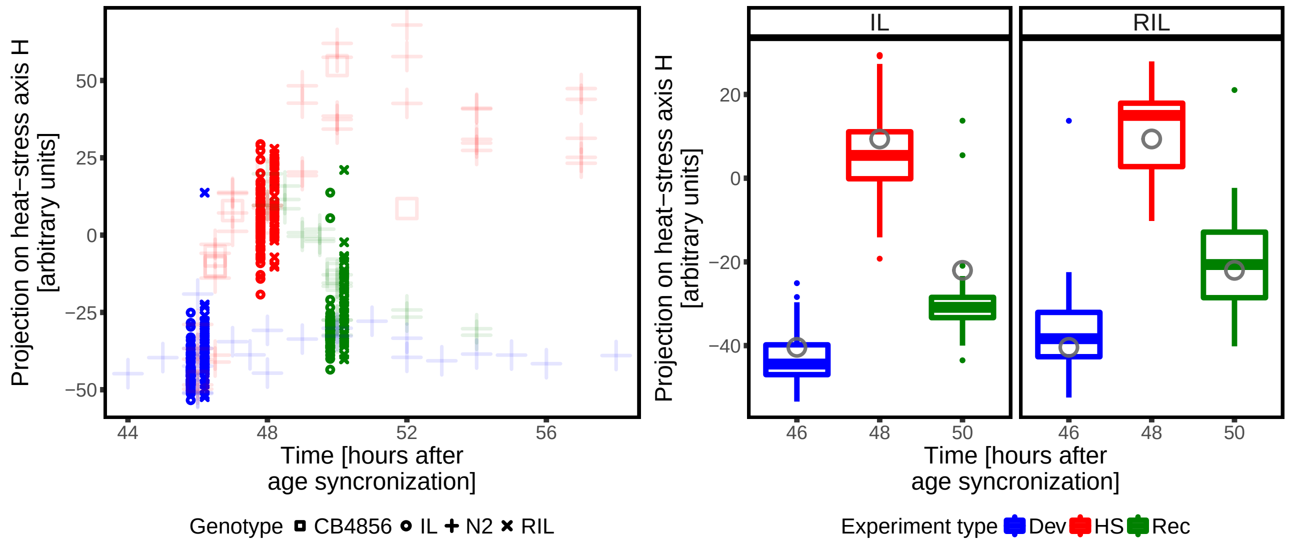


Supplementary Figure S8: Projection of the RILs and ILs samples on the heat-stress axis H. Different colors correspond to different treatments (blue is for unperturbed worms, red for heat-stressed samples, and green for recovering worms), while symbols refer to genotypes. The left panel shows the projection of all the data (data points of parental lines are in the background). The rights panel directly compare the projections of the ILs and RILs on the heat-stress axis under different treatments with the corresponding projection of N2 (averaged over replicates). The value of the projection for N2 is comparable with the projection for the RILs and ILs.

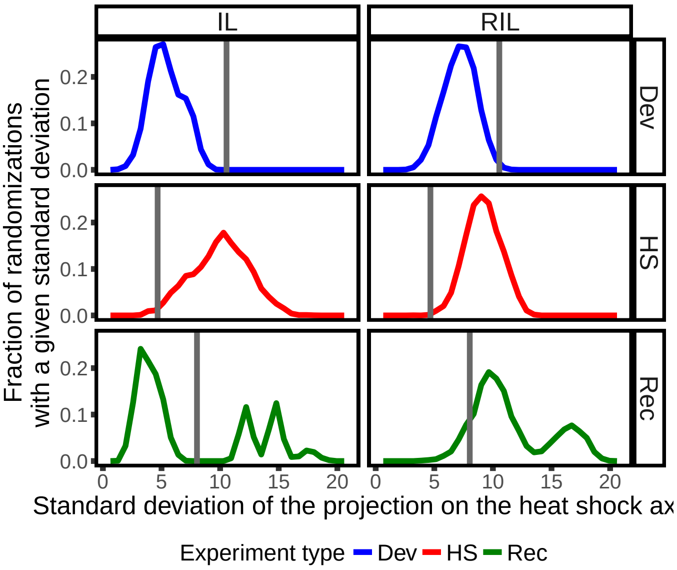


Supplementary Figure S9: Comparison between the variability of the projection on the heat-stress axis of RILs, ILs and N2. For each condition, we considered the replicates obtained with the N2 strain and measure the standard deviation of their projection across replicates (vertical gray bar). These values quantify the variability in the projection under different conditions. For each treatment, we then draw at random a number of RILs (or ILs) equivalent to the number of replicates available for N2 and measure the standard deviation of the random sample. By repeating the sampling 5000 times, we obtain a distribution of standard deviations (colored lines). The overlap between these distributions (colored lines) and the variability across replicates of the N2 strain quantifies whether genetic differences are responsible for the variability between RILs and ILs. The variability of RILs and ILs is comparable, resulting slightly larger in the RILs for the recovery experiment. In the case of unperturbed worms (blue line), the variability within N2 is larger than the variability between RILs. The projection obtained during heat-stress (red line) is instead more variable for RILs and ILs. The distribution of standard deviation for the random sampling of the recovery experiment (green line) is bimodal, due to the presence of outliers in the RILs and ILs, and their variability is comparable with the variability of N2.

In order to explore the possible sources of the variability between RILs and ILs in their projections, we compared it with the variability observed for N2 across replicates of the same experiment. For each experiment, we computed the standard deviation of the projection on the heat-stress axis of N2 with the one obtained by sampling at random an equal number of RILs or ILs. Figure S9 shows that the variability of the ILs and RILs is higher, lower or comparable than the one of N2, depending on the treatment. This suggests that there is not a detectable genetic signal on the projection on the heat-stress axis, and that the variability of the projection is most likely due to variability in the conditions and noise rather than biological differences.

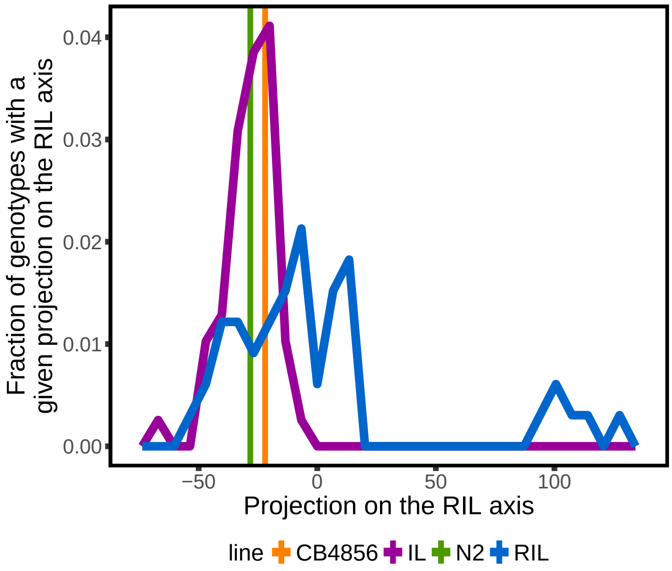


Supplementary Figure S10: Projection of RILs and ILs data on the RIL axis. The two vertical bars represent the projection of CB4856 (orange) and N2 (purple), averaged over replicates. The two lines are the distribution of the projections of RILs (blue line) and ILs (green line) on the RILs axis. The RILs axis was obtained as the second principal axis from the RILs data at 48h. All the worms considered in this figures were collected after growing under control conditions for 48h at 20°C degrees.

##### B. Derivation of the genetic heat-stress axis, GH

As shown in section S2 C the variability of the projection on the heat-stress axis between RILs and ILs is not particularly different from the variability observed across replicates of the N2 strains (see Figure S9). This is not surprising, given the method used to infer the heat-stress axis. This axis was in fact inferred using only the N2 strains under different exposure to heat-stress. It is therefore reasonable to think that both the variability between genotypes at the same time point is due to fluctuations of the conditions or other sources of noise and not due to biological differences between organisms.

In order to explore the intrinsic differences between different genotypes, we applied a similar method to the one explained in section S2 B. Using only the RILs data taken after 48h of unperturbed development (47 data points) we performed a PCA and isolated the second principal axis. This axis (RILs axis in the following) is the one that explains most of the variation between RILs at 48h in control conditions (20°C). Figure S10 shows the distribution of projections on this axis of RILs (used to infer the axis), but also of ILs and the two parental lines (N2 and CB4856). As expected, ILs show lower variability in the projection on this axis.

In order to measure the difference in the response to perturbations between different RILs, we then used the data obtained after a two hours heat-stress to obtain the principal axes. The second principal axis is the one containing the information on the different responses to heat-stress. It is also related to other differences between RILs and not

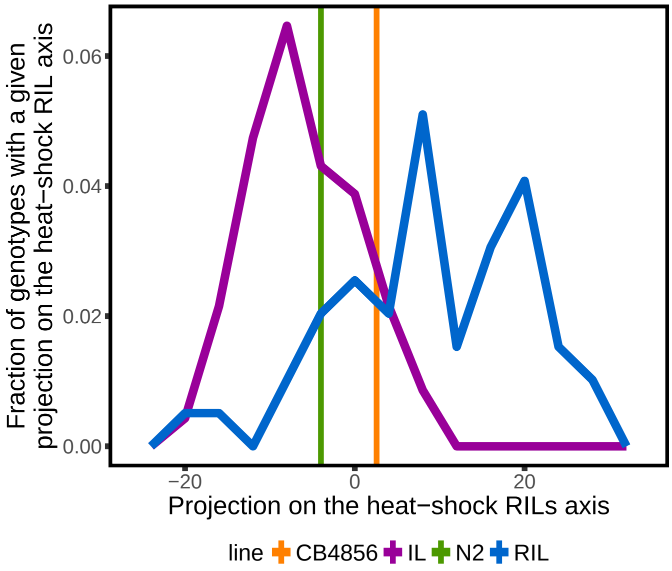


Supplementary Figure S11: Projection of RILs and ILs data on the RIL heat-stress axis, GH. The two vertical bars represent the projection of CB4856 (orange) and N2 (purple), averaged over replicates. The two lines are the distribution of the projections of RILs (blue line) and ILs (green line) on the RILs axis. The RILs axis was obtained as the second principal axis from the RILs data at 48h. All the worms considered in this figures were collected after 48h and grown at 20 degrees.

exclusively to their response to heat-stress. These other differences are the ones contributing to the RIL axis obtained using the expression before heat-stress. We obtained therefore a RILs heat-stress axis by removing the RILs axis from the second principal axis obtained with the heat-stress data (see section S2 B).

#### S4. LIFESPAN DATA

The lifespan data were collected independently from the data used to infer the axis. For each worm, survival was assessed every day until death (see Methods). The data were collected for different strains (the two parental lines, CB4856 and N2, the RILs and the ILs) and for two different condition: unperturbed ($20°C$ throughout life) and heat-stress (heat-stress perturbation at $35°C$ for 4 hours, see Methods). For each strain and treatment condition, the lifespans of an average number of 31 animals were scored.

An efficient way to analyze differences in lifespan consist in studying the differences between survival probabilities. At a given time $t$, one can measure the fraction of worms that are still alive $P_a(t)$. This quantity is simply related to the lifespan distribution $p(t)$ (i.e., the probability of dying exactly at time t), via

$$P_a(t) = \int_t^{\infty} ds\ p(s) \tag{9}$$

Figures S12 shows the survival probabilities $P_a(t)$ for different strains and the unperturbed condition, while figure S13 shows the same quantity for the heat-stress data. There is an observable variability between strains and

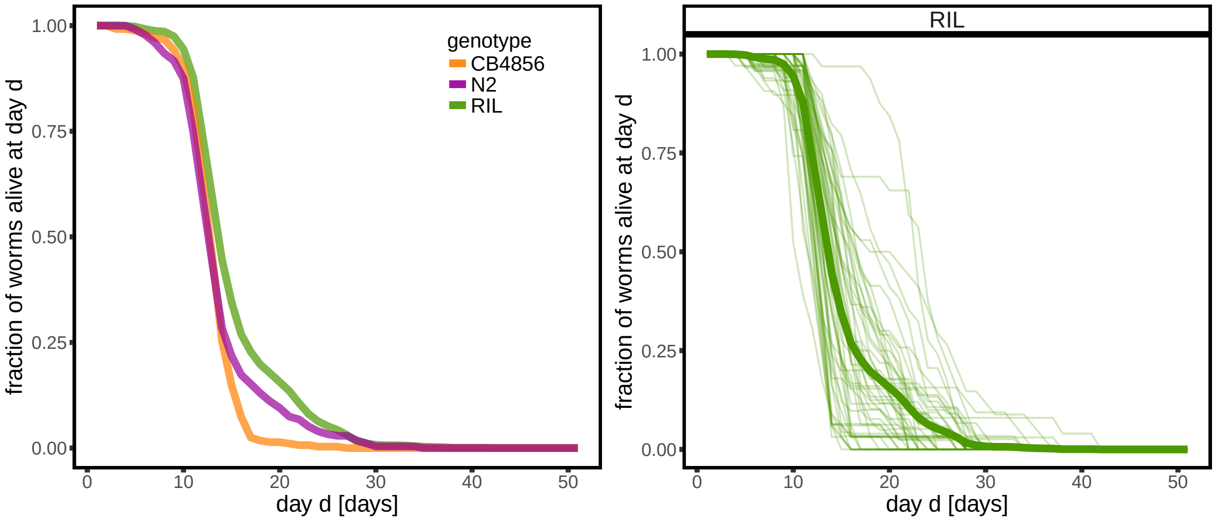


Supplementary Figure S12: Fraction of worms alive at a given day. The left panel show the survival curve for the two parental lines (N2 and CB4856) and the average across RILs and ILs. The two panels on the right shows the survival curves for each RIL and IL strain (thin lines) compared with the averages (thick line, also shown in the right panel). As expected and previously reported, the difference between different RILs (and ILs) in lifespan is much larger than the difference observed between N2 and CB4856.

between conditions. The shape of the survival curve is fairly similar among different strains, while it differs between the two conditions. It is important to observe that the variability between RILs (or ILs) is larger than the variability between the two parental lines.

As expected, the average lifespan is reduced when the worms are exposed to 4 hours of heat-stress (see Figure S14). Remarkably, the average lifespan with and without heat-stress are not correlated.

Figure S15 shows the lifespan distribution $p(t)$ for the two parental lines with and without heat-stress. In the case of the perturbed data, the distribution is bimodal, with a strong peak just after the heat-stress. This peak suggests that a non-negligible fraction of the worms dies because of the heat-stress, while other survive and their lifespan is affected by the heat-stress. Notably, the average lifespan of the perturbed worms is strongly anti-correlated with the survival in the first five days after the heat-stress (see figure S16). This correlation implies that the average lifespan for the unperturbed data should not be consider as the typical lifespan, but as related to the survival after the first days following heat-stress.

#### S5. HEAT-STRESS RESPONSE AND LIFESPAN

We correlated the projection on different axis of different treatments with the effect of the heat-stress on lifespan. To measure the effect of heat-stress on lifespan, we considered the ratio between the lifespan with and without heat-stress.

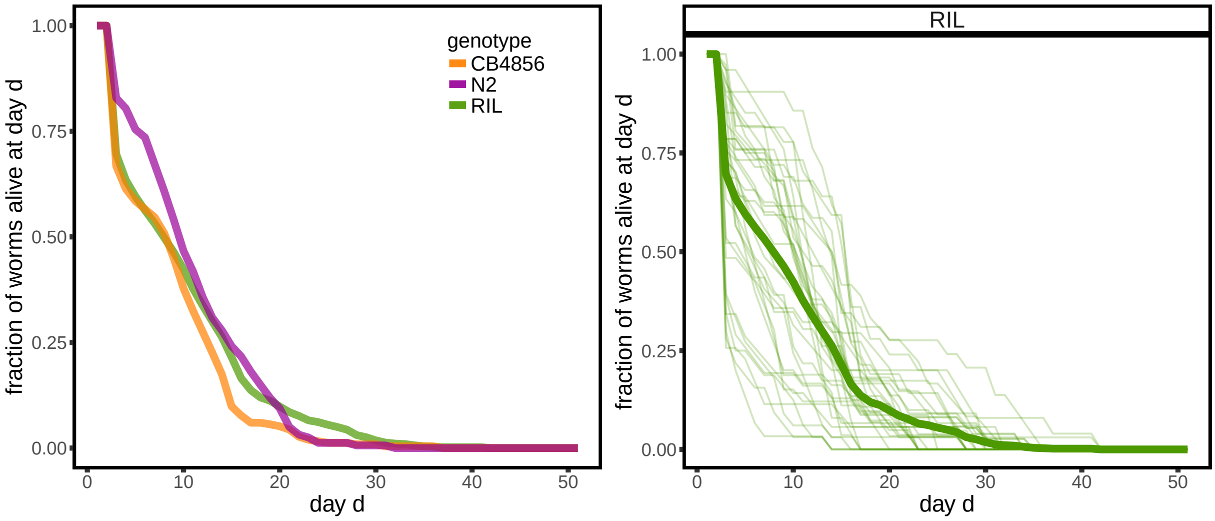


Supplementary Figure S13: Fraction of worms alive at a given age, after exposure to heat-stress at day 2. All the worms were exposed to a 4 hours heat-stress when they were 48 hours old (see Methods). The left panel show the survival curve for the two parental lines (N2 and CB4856) and the average across RILs and ILs. The two panels on the right shows the survival curves for each RIL and IL strain (thin lines) compared with the averages (thick line, also shown in the right panel). As expected and previously reported, the difference between different RILs (and ILs) in lifespan is much larger than the difference observed between N2 and CB4856.

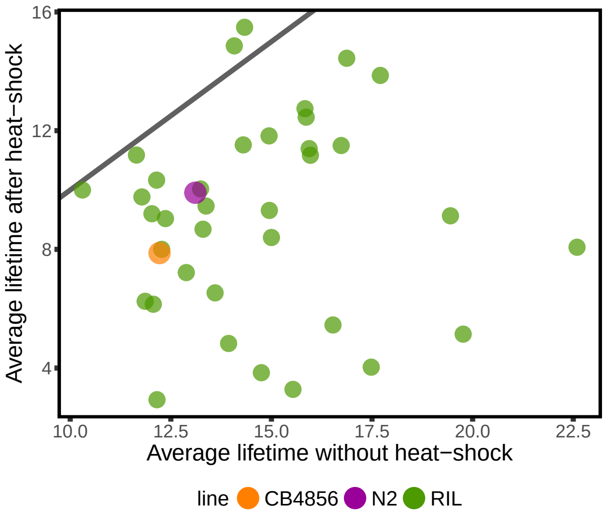


Supplementary Figure S14: Average lifespan after heat-stress vs. average lifespan without heat-stress for different strains. The continuous line is the identity line. As expected, the lifespan is reduced by heat-stress (*i.e.* the points tend to stay below the continuous line). On the other hand, there is no correlation between the two, implying that the effect of heat-stress on lifespan is not proportional to the average lifespan under control conditions.

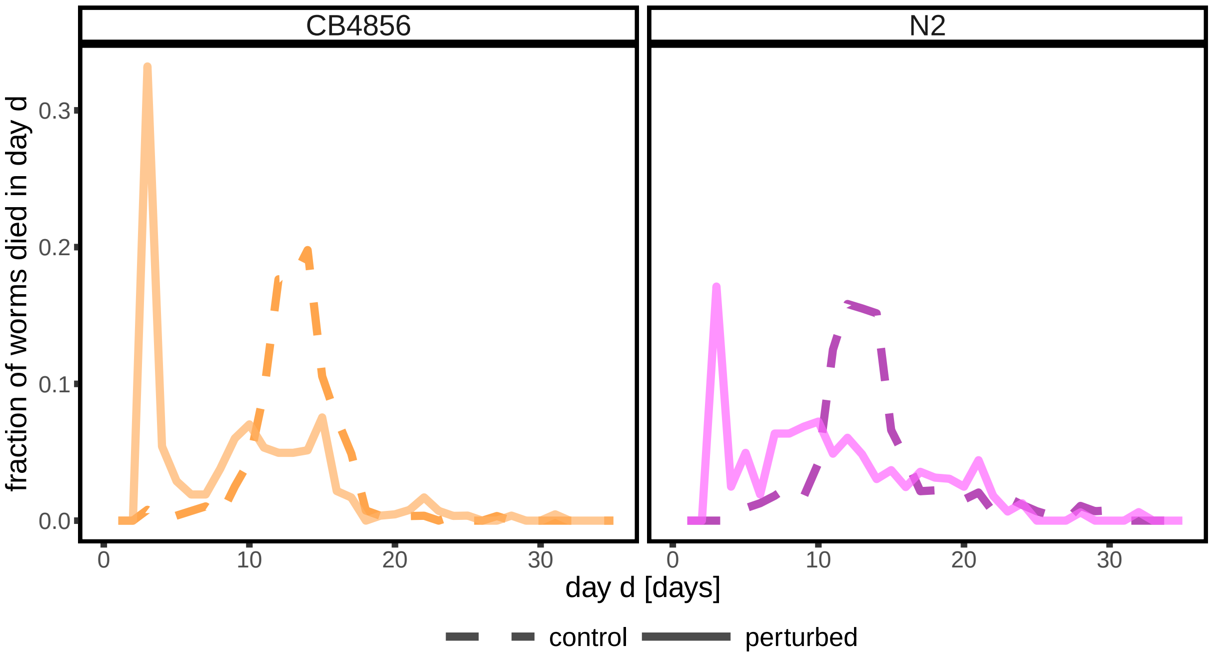


Supplementary Figure S15: Lifespan distribution of parental strains with and without heat-stress. Dashed lines were obtained from worms kept at $20°C$ for their entire life, while continuous lines are lifespan distributions of worms that were exposed to $35°C$ for 4 hours on the second day of their life. The lifespan distribution of the worms exposed to heat-stress shows a peak just after the exposure to heat-stress, suggesting the a significant fraction of worms dies just after the heat-stress.

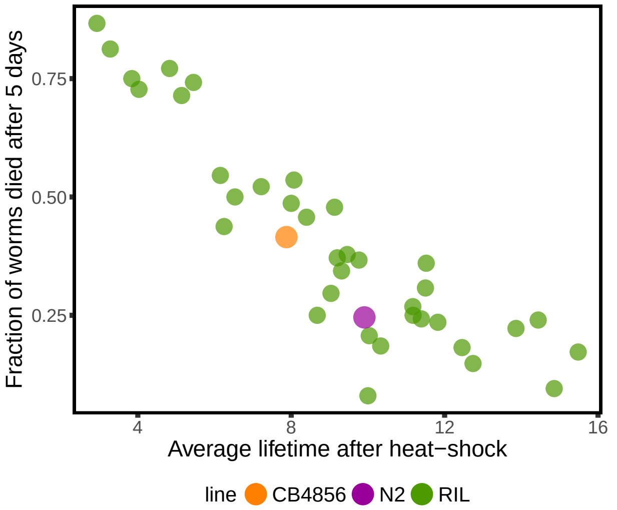


Supplementary Figure S16: Average lifespan after heat-stress vs fraction of worms dead before the 5th day. There is a strong anti-correlation, that suggests that the average lifespan in the presence of heat-stress is mostly determined by the first peak in figure S15.

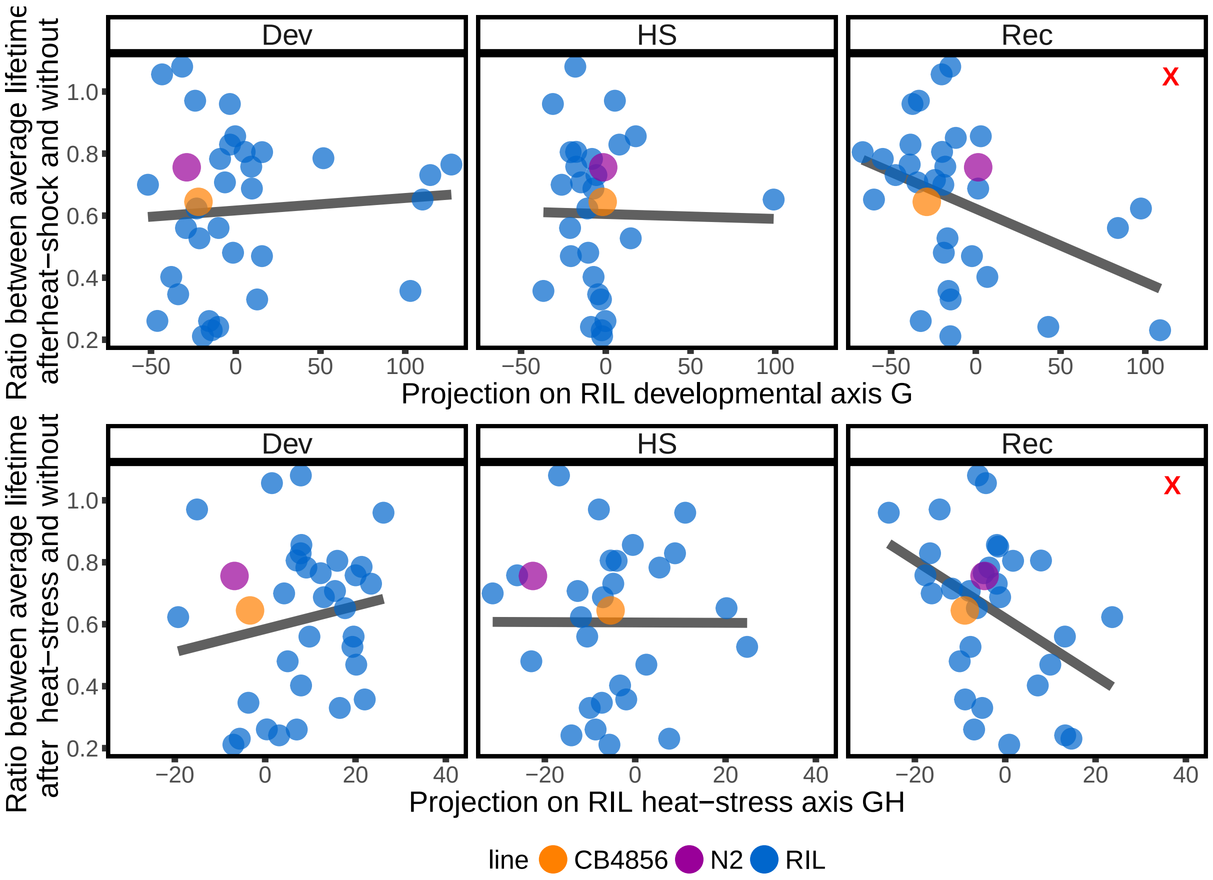


Supplementary Figure S17: Relation between the effect of heat-stress on lifespan and projection on different axis for RILs data. Each plot shows the correlation between the ratio of lifespan with and without heat-stress and the projection of a given condition on one axis. The upper panel shows the projections on the RIL axis (which encodes differences between RILs), while the lower panel shows the projections on the heat-stress RIL axis (which represents differences between heat-stress response of different RILs). The projections of the recovery data on both axes correlates with the effect on lifespan.

Figure S17 shows that the projection on these axis during recovery correlates with the effect of lifespan.

We used the same axes inferred by using the RILs data, to test whether the same correlation was present in the ILs. We found that the ratio between lifespan with and without heat-stress correlates with the projection of the recovery data on the heat-stress axis S18. We also found a correlation with the projection of the heat-stress data on the same axis that was not present in the RILs data.

**References**

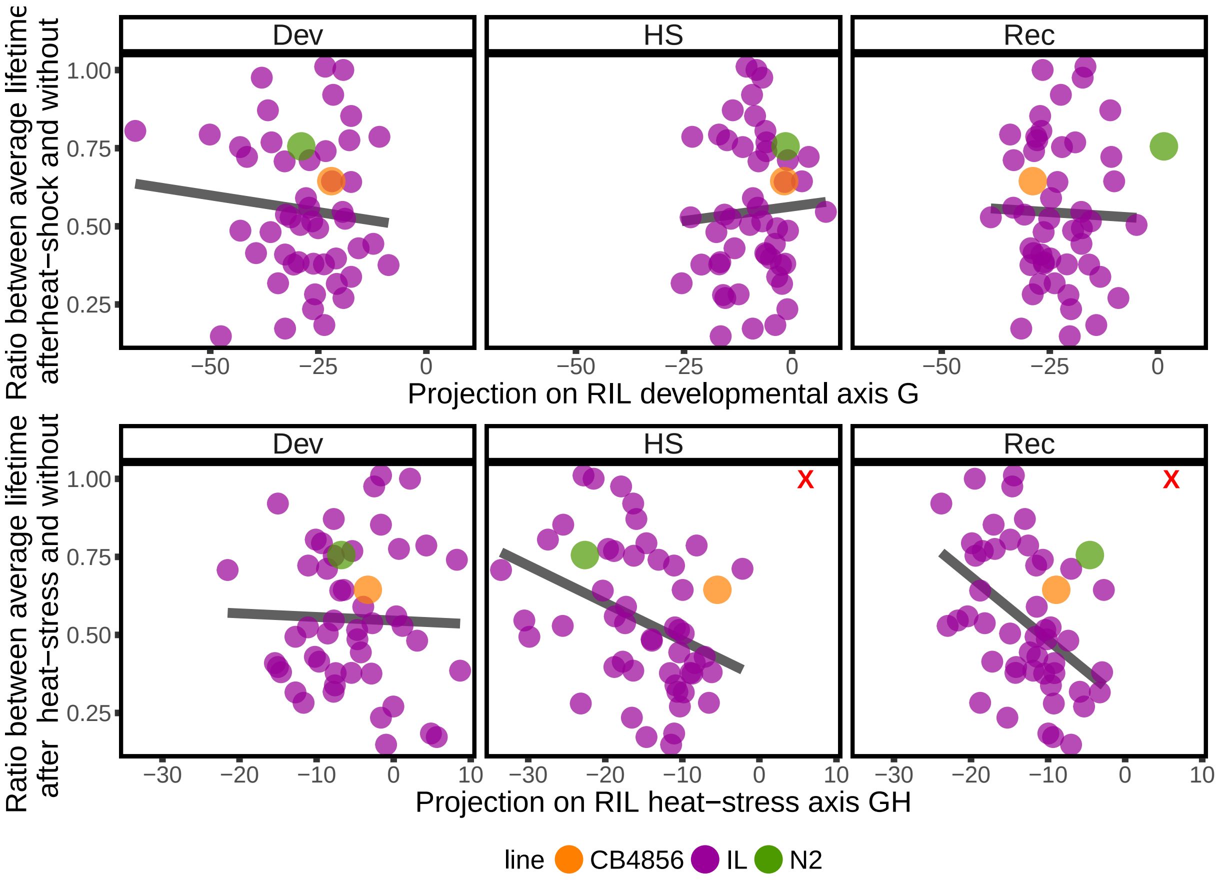


Supplementary Figure S18: Relation between the effect of heat-stress on lifespan and projection on different axis for ILs data. Each plot shows the correlation between the ratio of lifespan with and without heat-stress and the projection of a given condition on one axis. The upper panel shows the projections on the RIL axis (which encodes differences between RILs), while the lower panel shows the projections on the heat-stress RIL axis (which represents differences between heat-stress response of different RILs). The projections of the recovery data and the heat-stress data on the RIL heat-stress axis correlate with the effect on lifespan. Note that the axes were obtained independently of this dataset using only the RILs data.
